# Isolating and Quantifying the Role of Developmental Noise in Generating Phenotypic Variation

**DOI:** 10.1101/334961

**Authors:** Maria Kiskowski, Tilmann Glimm, Nickolas Moreno, Tony Gamble, Ylenia Chiari

## Abstract

Phenotypic variation in organisms is typically attributed to genotypic variation, environmental variation, and their interaction. Developmental noise, which arises from stochasticity in cellular and molecular processes occurring during development when genotype and environment are fixed, also contributes to phenotypic variation. The potential influence of developmental noise is likely underestimated in studies of phenotypic variation due to intrinsic mechanisms within organisms that stabilize phenotypes and decrease variation. Since we are just beginning to appreciate the extent to which phenotypic variation due to stochasticity is potentially adaptive, the contribution of developmental noise to phenotypic variation must be separated and measured to fully understand its role in evolution. Here, we show that phenotypic variation due to genotype and environment, versus the contribution of developmental noise, can be distinguished for leopard gecko (*Eublepharis macularius*) head color patterns using mathematical simulations that model the role of random variation (corresponding to developmental noise) in patterning. Specifically, we modified the parameters of simulations corresponding to genetic and environmental variation to generate the full range of phenotypic variation in color pattern seen on the heads of eight leopard geckos. We observed that over the range of these parameters, the component of variation due to genotype and environment exceeds that due to developmental noise in the studied gecko cohort. However, the effect of developmental noise on patterning is also substantial. This approach can be applied to any regular morphological trait that results from self-organized processes such as reaction-diffusion mechanisms, including the frequently found striped and spotted patterns of animal pigmentation patterning, patterning of bones in vertebrate limbs, body segmentation in segmented animals. Our approach addresses one of the major goals of evolutionary biology: to define the role of stochasticity in shaping phenotypic variation.

## 1. Introduction

A first-order approximation of phenotypic variation is that genotype and environment variation maps to phenotype variation. Stochastic variation is an important third source of phenotype variation included in a more complete description of the genotype-phenotype mapping [1-4]. Choosing among a variety of terms found in the literature, phenotypic *variation* is the diversity in a trait attributed to genotypic variation (differences in genotype), phenotypic *plasticity* (differences in environment), and *developmental noise* (the difference in phenotypic outcomes that occurs when genotype and environment are fixed (e.g.,[5-7]), due to stochasticity in cellular and molecular processes [4, 6]). The interaction of these sources of variation, combined with the sheer magnitude of the number of phenotypic variables, makes the study of the phenotypic variation of a population extremely complex.

Computational morphometric approaches may identify a small number of key phenotype features (comparable “units” [8]) or apply clustering analyses over a higher number of phenotype variables, to objectively sort complex phenotypic variations and quantify differences between them. Here we describe a computational approach for interpreting phenotype differences in the context of the contribution of genotype, environment, and developmental noise, by applying a computational model of the organismal development of these phenotypes. In our stochastic simulations, fixed (predetermined) parameters correspond to genetic and environmental factors while intrinsic stochastic variation within simulations corresponds to development noise. By applying a computational model of a developmental mechanism to generate phenotypes, such as the reaction diffusion model we use here, the simulated phenotype variation will reflect the intrinsic freedoms and constraints of the developmental mechanism itself [9, 10].

### Periodic Color Patterning in Vertebrates: a likely developmental mechanism

Vertebrates show a wide variety of integumentary colors and patterns both within and among species. Variation in vertebrate coloration represents a model to understand the link between genetic basis, developmental patterns, and phenotype (*e.g*., [11-14]). Furthermore, variation in coloration is often under strong selection and linked to ecological or behavioral differences within and among species (*e.g*., [9, 15-18]). Although the genetic and developmental basis of vertebrate coloration have been identified for a few species (*e.g*., [11, 14, 19]), the mechanisms involved in color pattern formation in vertebrates are largely unknown (but see for example [20] and references therein; [21]), especially for vertebrates other than mammals or fish. While molecular approaches can uncover the genes involved in determining a certain color pattern (e.g., [19]), most of the time, especially for non-model species, the relationship between the candidate genes involved in color pattern formation, organization and variation and the observed phenotype are unidentified [14].

Strikingly, body color patterns with periodicity such as spots and stripes are found ubiquitously throughout vertebrates (studied most extensively among cats, fish, and some reptiles, see below) suggesting that mechanisms for periodic patterning may be very common, even universal, among vertebrates and thus their development conserved among organisms. The mechanism of ‘local activation long range inhibition’ (LALI; [22-24]) represents a general theoretical model predicting patterns that are spotted, striped or of an intermediate mixed form (‘labyrinthine’). The most important such mechanism is the Turing mechanism in reaction-diffusion system, where the local activation and lateral inhibition are due to reaction kinetics mediated by diffusion [25]. For mammalian coat pattern formation, Murray was the first to propose an activator-inhibitor LALI mechanism [26-28], with the idea that a chemical pre-pattern established by a Turing-type mechanism dictates cell differentiation. Murray showed that many mammalian patterns observed in nature can be produced by such a mechanism. Turing-type mechanisms have since become a frequently studied and widely hypothesized mechanism for periodic patterning of integument in vertebrates (a wide array of mammals, including cats [15], several species of fish [21, 29-31] and reptiles, especially squamates such as snakes [9, 15, 32, 33] and recently a convincing ‘living’ (experimental) reaction diffusion model for skin color patterns in the ocellated lizard [34].

Two key characteristics of patterns generated by LALI mechanisms are: 1) non-random regularity in the spacing of clustered elements (periodicity) and 2) potential for mixed transitions between spotted and striped patterns. Thus, the LALI model is suited to analyze periodic patterns and can provide a mathematical framework for analyzing pattern transitions between spotted and striped phenotypes. Many species of lizard and snake (squamate reptiles) demonstrate all of these pattern variations during ontogeny, within the same individual among the different parts (*e.g*., tail, trunk and head), and among individuals of the same or different species (YC pers. obs.). Among reptiles, the leopard gecko (*Eublepharis macularius*) shows particularly dramatic changes in pattern during maturation (Figure 1), transitioning from a hatchling pattern with alternating dark and light bands with a dark head to an adult pattern consisting of a light colored body and head with scattered dark spots [35, 36]. Leopard geckos have been bred in captivity for decades and during that time numerous color and pattern mutations have been developed by private hobbyists [37, 38], providing a unique opportunity to understand how pattern variation can be created at the intraspecific level.

**Figure 1:**
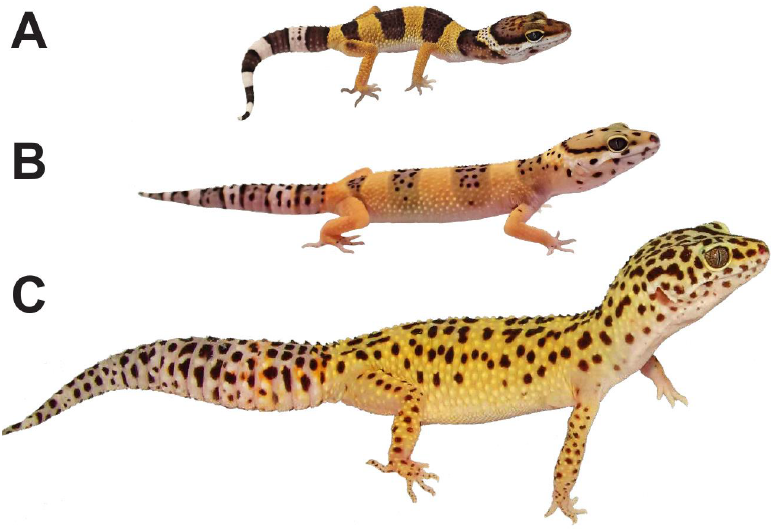
Ontogenetic pattern change in the leopard gecko (*Eublepharis macularius*). Each photographed individual represents the typical color and pattern of a (A) hatchling (one month old), (B) juvenile (three months old), and (C) adult (>12 months old) gecko. Relative sizes are approximate and the hatchling image has been enlarged to allow an easier comparison of pattern detail among individuals. Pictures by T. Gamble.

Since the leopard gecko demonstrates such a broad range of stage and body-plan-specific patterning, in this work, we focus on understanding the mechanisms generating the color pattern variation of a precise region on the leopard gecko head (the parietal, post-orbital region) during a specific stage of their development (at nine weeks). Among all the individuals analyzed in this study, this region of the head at nine weeks is invariably a simple spot pattern of discrete melanistic blotches on a pale background. By analyzing images of these regions, we extracted key morphological features of the spotted pattern. These extracted features allowed us to compare the gecko’s patterns with patterns obtained through LALI simulations: a simulation pattern and a gecko pattern were defined to be “matching” if they had the same values for these morphological features (see below). We were able to find matches by varying LALI parameters (corresponding to changing genetic and environmental factors) for the eight experimentally observed gecko patterns within a low-dimensional LALI space (see definition in Table 1). By varying parameters within this region of LALI space, we were able to generate very likely, “normal” pattern variations. Furthermore, by expanding parameters just outside this region, we were able to generate “preternatural” pattern variations predicted by the developmental model just beyond the region of natural variation. These preternatural patterns are patterns that are not observed on the studied geckos, but are patterns that could potentially exist by varying the genotype and environment. These preternatural patterns in fact look like typical periodic patterns that are seen on animals, but they have morphological features with values that are slightly lower or higher than those we observed on the eight studied geckos.

**Table 1:** Definitions of the terms used in this manuscript relating to the *L*→*P* map.

|  |  |
| --- | --- |
| LALI mechanism | A LALI mechanism is a mechanism for periodic patterning based on local activation and long range inhibition (LALI; [22-24]). |
| LALI-type | The LALI-type is the set of parameters needed to specify a LALI mechanism, for either a computational or a natural biological process. As parameters are varied systematically in the computational model, these result in predictable pattern differences such as smaller or larger spots, smaller or larger spot separations, or spot to stripe transitions. (This is analogous to the combined effects of genotype and environment, but excludes developmental noise.) |
| LALI-space | The set of all possible LALI-types. |
| Phenotype | A set of measured observables (e.g. morphometric quantities or “comparable units” (Kuhl and Burghardt, 2013), selected from a larger set of all the possible observables that could be measured from a complex pattern or organism. |
| Phenotype Space | The set of all possible phenotypes, for the selected set of phenotypes (measured observables). |
| $L \rightarrow P$ Map | The LALI-type to phenotype map is the observed mapping, either computational or biological, from a set of LALI-types to a set of phenotypes. Due to random variation, one LALI-type can map to a set of phenotypes (see ‘phenotype cloud’), and many LALI-types can map to one phenotype (see ‘neutral region’). |
| Phenotype Cloud of a LALI-type | Due to random variation (corresponding to developmental noise), a set of LALI parameters can result in many different phenotypes. This set of phenotypes is the ‘phenotype cloud’ of that LALI-type. (This is roughly analogous to the concept of reaction norm in ecology and evolution.) |
| Neutral Region of a Phenotype | A phenotype can be produced by many LALI-types. The neutral region of a phenotype is the set of LALI-types that are likely to yield that phenotype. (This is analogous to the genomic neutral region of a phenotype.) We define the $r\%$ neutral region of a phenotype $P$ as the set of all LALI-types $L$ such that the $r\%$ phenotype cloud of the LALI-type includes the phenotype $P$. |

### LALI-type to Phenotype Map *L→P*

Many mechanisms with the same core LALI logic (molecular, cell-based and/or mechanical) yield similar patterning despite different underlying biological processes [39]. While variations of reaction diffusion process are often used to explain Turing patterns, other candidate mechanisms include cell-based and mechanical processes [39]. In vivo, it has recently been established that Turing patterns on zebrafish skin are the result of a mechanism that satisfies the core LALI logic but that is qualitatively different from reaction diffusion [40]. With this perspective in mind, especially given that the molecular details of leopard gecko skin patterning remains unknown, our aim was to investigate mathematical features of LALI mechanisms in general rather than commit to a specific one. We therefore compared and contrasted results for two computational models – a model based on linear reaction dynamics (as in Turing’s classic paper [25]), and one based on FitzHugh-Nagumo reaction kinetics [41-43].

In biological systems, a LALI mechanism depends on physical quantities such as molecular diffusion rates, protein reaction rates, cell response rates, material resistance to bending and compression, etc. Computational LALI models aim to simulate the *net* effect of these physical quantities with a relatively small number of parameters. For example, the operator ℒ of the Swift-Hohenberg equation [39] summarizes the net outcome of non-local effects. By simulating the net effect of these physical quantities, pattern variation can be investigated without needing to fit a large number of unknown physical quantities. Given either a computational model of a LALI mechanism or a real world LALI mechanism (e.g., color patterning), the LALI-type (Table 1 for definition) to phenotype map *L → p* is the map from parameters (see below) of the LALI model (‘L’) to a final phenotype (‘P’). It represents a developmental step that occurs after genotype and environment have determined the parameters relevant to the LALI mechanism (see Figure 2), for example after the environment and genotype determines which gene(s) should be activated, how much, and for how long in a certain body region. Varying the parameters of the LALI-type in simulations is analogous to varying the underlying genetic and environmental factors that specify the LALI mechanism (e.g., molecular, cell-based, mechanical) producing a certain phenotype in biological systems.

**Figure 2:**
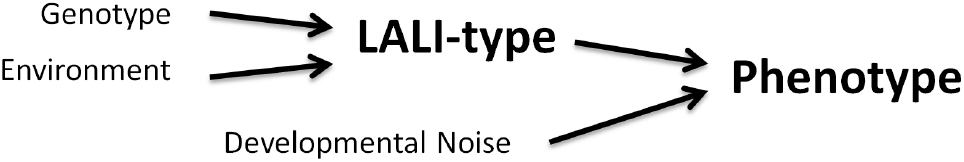
Conceptual Model of the LALI-type to Phenotype Map. The LALI-type summarizes the genetic and environmental factors of a LALI pattern. The phenotype is a product of the LALI-type and stochastic effects (random variation called developmental noise).

Furthermore, given either a computational model or an actual biological LALI mechanism, the spatiotemporal evolution of the pattern is influenced by developmental noise. Indeed, the diffusion-driven instability in both the linear model and the FitzHugh-Nagumo model relies on small, stochastic fluctuations of the micro-environment, just like any Turing-like instability in pattern formation mechanisms with a core LALI logic [39]. The diffusion-driven instability occurs through the amplification of these fluctuations and ultimately creates the Turing pattern of regions (e.g., spots or stripes) of high and low concentrations of some chemical (“peaks and troughs” [25]; or Turing “bifurcation” [40]). Such stochastic fluctuations enter the model through randomly perturbed initial conditions. The effect of stochastic variation in the computational LALI model is that the relationship between LALI-type and phenotype is not one-to-one, that is, even if genetic and environmental factors are fixed, a single LALI-type can generate different phenotypes and conversely different LALI-types may generate the same phenotype. Thus, this framework and a computational model of a LALI mechanism permit to either vary genetic and environmental factors or hold them fixed (by varying or fixing the parameters of the computational LALI model), and to investigate the outcome of the *L*→*P* map with respect to different factors contributing to it (environment, genotype, and developmental noise). Specifically, the approach used in this work allows isolating the effect of genetic and environmental variation versus random variation and specifically to quantify the role of developmental noise on the *L*→*P* map.

## 2. Methods

### A. Pattern Analysis and Morphometrics of Live Geckos

#### Data collection

Individual live leopard geckos were photographed within two weeks of hatching and subsequently photographed every two weeks thereafter to document pattern changes over time. Each gecko was photographed on a smooth surface that featured a grid pattern with evenly spaced lines 12.7 mm apart. Photos were taken with a Sony DSC F828 8 megapixel camera mounted on a tripod approximately 45 cm above the gecko. One single photo per gecko was used in this study.

#### Pattern Selection

For pattern analysis, we selected a disk-shaped region of the parietal, post-orbital head region of eight geckos at nine weeks (see Introduction regarding selection of chosen developmental time). A disk-shaped region was identified from images of the gecko head by an algorithm run in Matlab (Matlab R2015b, Mathworks, Inc.) that identified the largest disk that could be inscribed within the boundary of the head, for a disk centered at the centroid of the head. The size of the pattern disk varied for each gecko, but was invariably a pattern of isolated melanistic spots on a lighter background. Images of the eight gecko heads and the corresponding selected disk-shaped regions are shown in the first and second columns of Figure 3. Focusing on this region restricted the image to one pattern type (discrete dark spots on a lighter background) in order to study a homologous region among individuals without including areas of the head where the pattern transitioned from one type of pattern to another. Head boundaries of the gecko were determined by hand, while identifying the centroid of the head and the largest inscribed disk was automated for the eight gecko images using Matlab.

**Figure 3:**
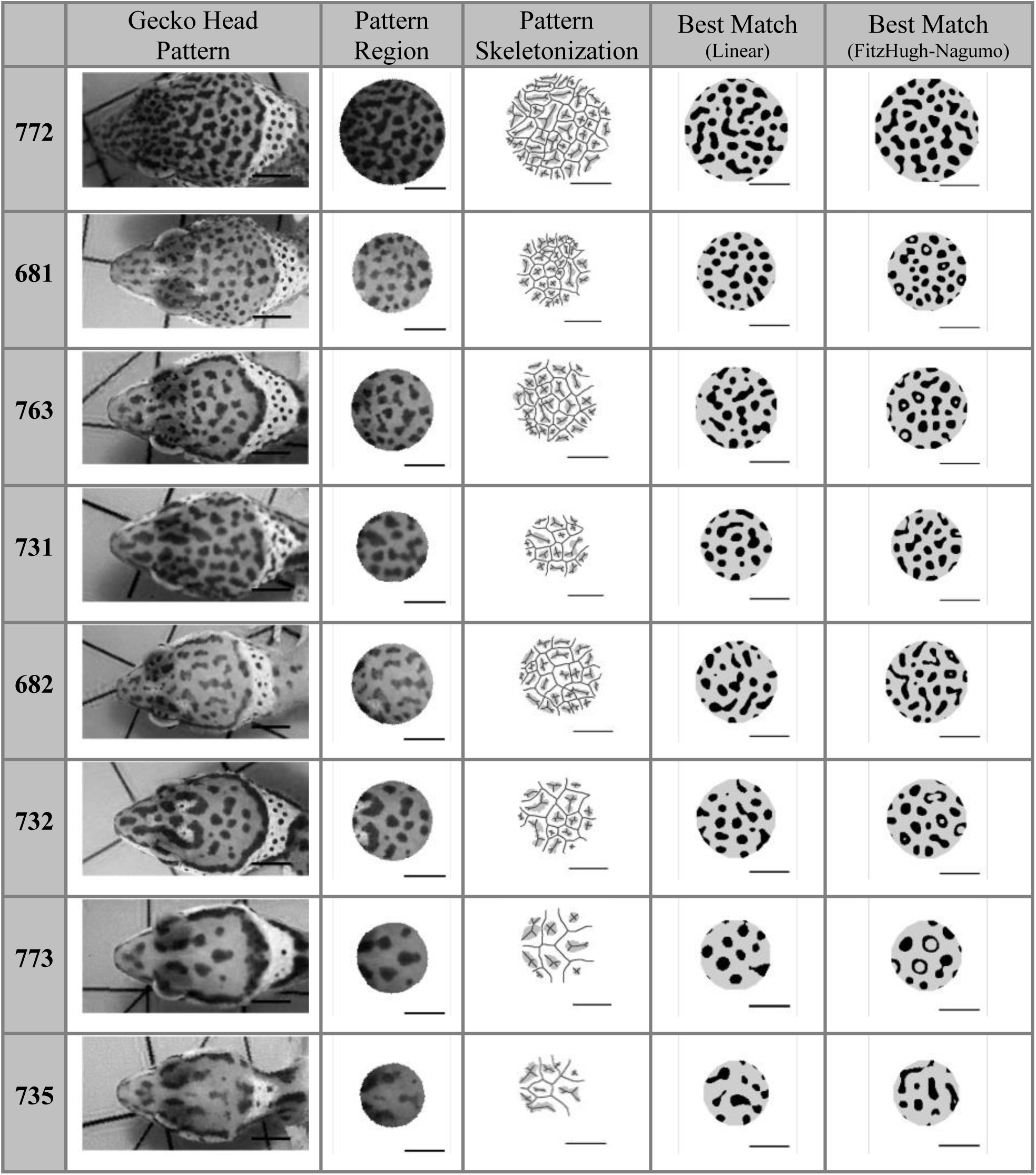
Automated disk-shaped pattern selection of parietal, post-orbital head region of eight geckos at nine weeks. For each Gecko ID (numbers on the left): **Left**: Images of the eight gecko heads at nine weeks; **Second column**: the disk-shaped parietal, post-orbital (DSPPO) region that was selected for pattern analysis, preserving their relative sizes; **Third column:** Final pigment pattern identified by image analysis with the skeletonization of the image overlaid. **Fourth column:** Best phenotype match of 100 patterns simulated by the corresponding LALI-type using the linear model. **Right**: Best phenotype match of 100 patterns simulated by the corresponding LALI-type using the FitzHugh-Nagumo model. Horizontal bars indicate 0.5 cm. Geckos are ordered by decreasing fractional spot area of the pattern (see Table 2 for definitions).

#### Image Processing

Image processing of the live gecko images was required to correct for uneven lighting within and between photos, as well as different background levels of pigment from one gecko to another. Each disk-shaped image was contrast-enhanced (using Matlab’s internal *adapthisteq* function) to correct for shadows and inconsistent lighting. A threshold was then applied to binarize the image into a set of black pixels on a white background. Since pigment levels varied for each gecko, a different threshold value of pigment was required to discriminate spots from non-spots. For each gecko, the threshold value was calculated from the average pixel intensity *μ* and standard deviation σ of the pixel intensity. With experimentation, it was determined that a threshold of T=(μ - σ) was high enough to detect spots yet also low enough to identify their separations (i.e. pixels were required to be a standard deviation darker than average in order to be identified as a spot pixel). Thus, while each image had a different threshold for discriminating spots from non-spots, a single, objectively set definition was used to define this threshold. It was observed that small changes in the threshold could result in the joining or separation of nearby spots. To describe the variation generated by small variations in the choice of threshold, the mean and standard deviation of pattern statistics were computed by varying the threshold by 25% of the standard deviation of the pixel intensity. Before computing final pattern statistics, pattern noise due to arbitrary threshold cut-offs was eliminated by 1) filling holes within spots and 2) deleting stray pixels outside of spots. Holes within spots were identified as pixels with intensity below the threshold that were nevertheless completely contained within a region of pixels identified as a spot. Stray pixels outside of spots for removal were identified by a total area that was too small to be identified as a spot (the cut-off was 10 photo pixels). For the calculation of the *peak length* (Table 2), we found that the image skeletonization generated by the native Matlab operation could be sensitive to the contour of spots. Since the peak length should only depend on the spacing of spots and not their contour, we first found the convex hull of the spots and then skeletonized the image.

**Table 2.**
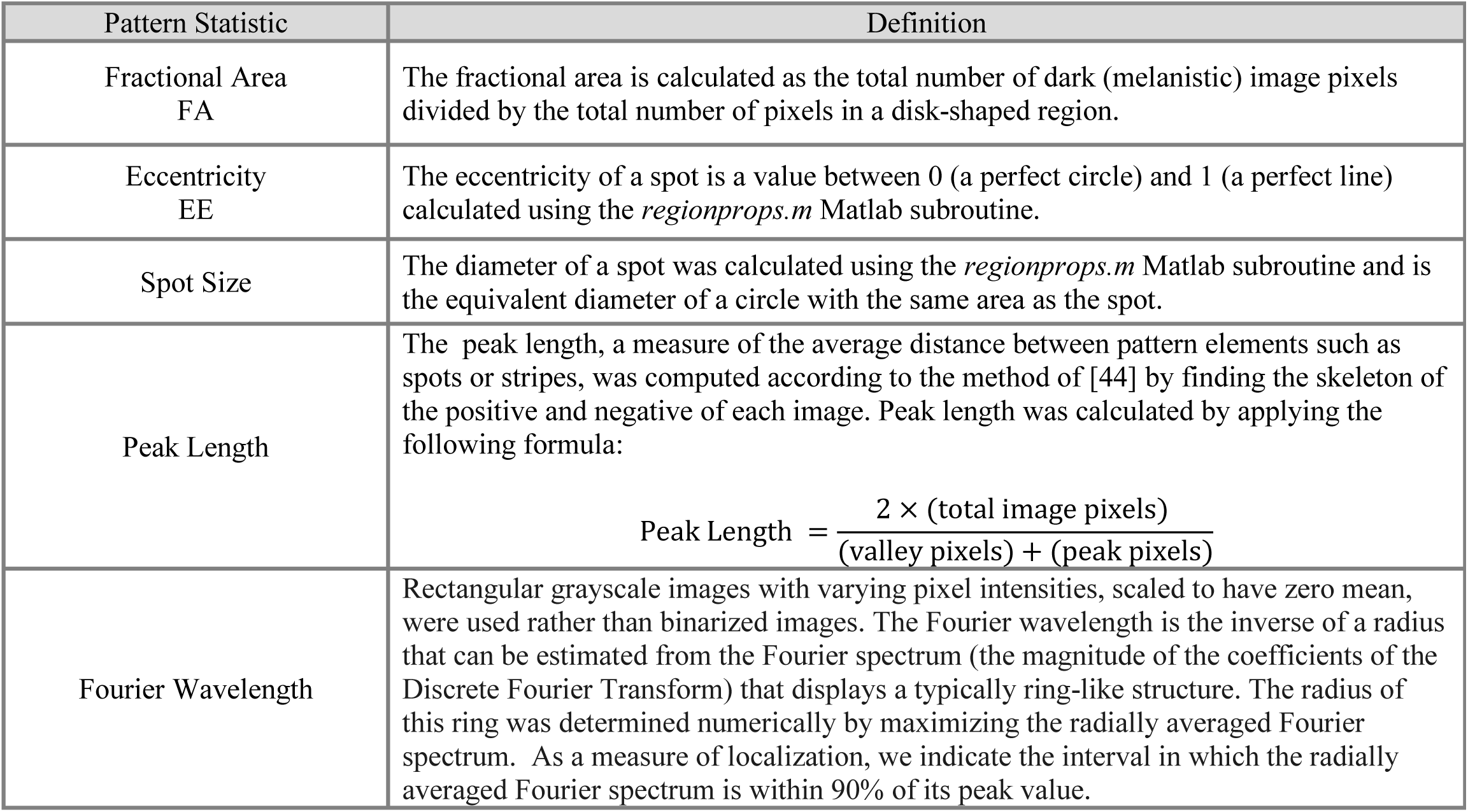
Morphometric properties and how they were calculated.

#### Morphometrics: Selected Phenotype Specified by Fractional Area and Eccentricity

For the pattern selected from the chosen region of the gecko’s head, we defined the *selected phenotype* of that pattern as the pair of numbers given by the *fractional area* and the *eccentricity* of the melanistic spots. The fractional area is a measure of the relative density of spots (pigmented regions) and the eccentricity of a spot is a measure of the pattern location on a spot to stripe continuum that is valid up until pigmented regions begin to overlap (see Table 2 for all definitions of technical terms used in this section). These two measures were chosen from an unlimited number of possible morphometric measurements to focus our analysis since these are measures that are readily interpreted in the context of periodic patterns (as measures of the relative spot or stripe size and position of the pattern along the spot to stripe transition), straight-forward to vary with LALI parameters (e.g., refs) and are scale free, and thus not depending on the size of the individual gecko. For both live and simulated pattern images, statistics were measured using automated Matlab scripts (these scripts accept as input any disk-shaped region with a binary pattern of black and white pixels, and did not distinguish between simulated and live gecko patterns). For the live gecko images, we also measured the average spot size, a scale-dependent statistic which was used for fixing the spatial scale of simulated pattern images (see discussion below of the selection for the pixel spatial scale). We also calculated the average distance between spots (i.e., the *peak length* or *Fourier wavelength* of the pattern).

### B. Simulation-Based Generation of LALI Patterns Using Two Reaction Diffusion Models

We modeled the core LALI logic of an activator-inhibitor system using two model implementations. Comparing results for two implementations helps identify which aspects of the LALI-space to phenotype space map may be most implementation specific. We used a linear Turing model implemented on a discrete cellular automaton and the non-linear FitzHugh-Nagumo model. In both models, *D* describes the relative diffusion rate of the activator *u* and inhibitor *v*.

*Linear Turing model*: The equations for the linear Turing model are given by:

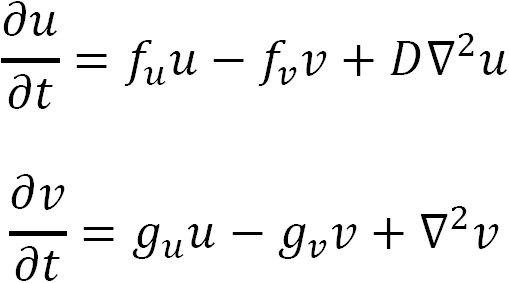

The parameters *f*_*v*_, *f*_*u*_, *g*_*u*_, *g*_*v*_ are fixed parameters that are initialized at the beginning of a simulation and do not change during the simulation. In an activator-inhibitor morphogen context, the parameters *f*_*u*_, *f*_*v*_, *g*_*u*_, *g*_*v*_ are understood, respectively, as the self-upregulation rate of the activator, the down-regulation rate of activator by inhibitor, the upregulation of inhibitor by activator and the self-down-regulation of inhibitor, respectively. Stochasticity is incorporated by initial conditions of random morphogen concentrations (initial concentrations of the activator *u* and inhibitor *v*) and stochasticity in the diffusion process (morphogens diffuse by random walk on a square grid). Simulations were run on a patterning domain consisting of a 200×200 spatial grid with periodic boundary conditions for a fixed number of time steps (200K). For all simulations for linear reaction diffusion in this manuscript, we found that it was sufficient to fix *f*_*v*_, **g**_*u*_, *g*_*v*_ and used the production rate of the activator *f*_*u*_ as a parameter for pattern matching (see also section C, step 2 of the Results).

*FitzHugh-Nagumo model*: The equations for the FitzHugh-Nagumo model are given by the following nonlinear reaction-diffusion equations:

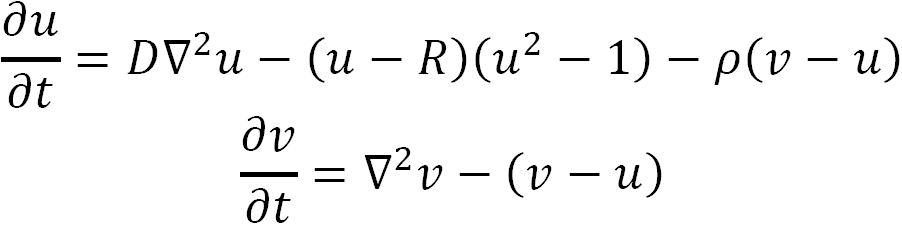

Here *D,R* and ρ are parameters, where *D* is the ratio of diffusion coefficients of the activator and the inhibitor, *ρ* is related to the relative rates of production of the activator and the inhibitor and *R* is a reference activator concentration. It is known that for certain ranges of parameters, the equations can produce stable spots or stable stripes in two dimensions [45]. We fixed *R* =0.047 and *D* =0.0194 and used *ρ* as a parameter for pattern matching. As a second parameter, we also varied the threshold value of morphogen that would map to whether a region was pigmented (see below). Stochasticity is incorporated by initial conditions of random morphogen concentrations. The equations were solved with the method of lines on square of side length *L*=10 with no-flux boundary conditions, using Matlab’s differential equation solver *ode45* (grid size of the discretization was 100×100).

For a complete list of model parameters, we also varied the threshold value of morphogen that would map to whether a region was pigmented (see below). Including this threshold parameter, the LALI-type for our linear model is completely specified by the five parameters [*f*_*u*_, *f*_*v*_, *g*_*u*_,*g*_*v*_,*T*] and the LALI-type for the FitzHugh-Nagumo model is completely specified by the four parameters [*D,R,ρ,T*].

*Final LALI Spot Patterns, Simulation Image Analysis and Pattern Statistics:* For both the linear and non-linear reaction diffusion models, simulations within an appropriate range of parameters resulted in a spatial pattern of low and high morphogen concentrations. Morphogen concentrations after a fixed number of simulation steps (200K steps for the linear model, time interval [0,20] for the FitzHugh-Nagumo model) were binarized into distinct regions of spots and non-spots using a threshold parameter *T* to produce a black and white pattern. In principle, the applied threshold parameter *T* in the computational models would correspond to the threshold concentration of morphogen for which biological cells produce pigment. The fractional area and eccentricity of the spots were calculated, for the entire simulated domain. Since fractional area and eccentricity are spatial-scale invariant, the generated *L*→*P* map that maps from model parameters to[*FA,EE*]-phenotype space is also scale-invariant and we consider that the absolute spatial scale of the pattern is a free parameter.

*Cutting Disk-Shaped Regions of Simulated Patterns*: For generating the simulated phenotype matches in Figure 3 and determining the intrinsic variability of each of the eight phenotypes in Figure 10, disks were cut from simulated domains to match the relative domain-to-pattern spatial scales of each of the eight phenotypes. First, since the spatial scale of a pixel in simulated patterns is a free parameter, the spatial size of a pixel was set so that the average size of spots in the simulated pattern would match those of the gecko phenotype. Once the spatial scale was established in this way, a disk was cut from the simulated pattern with the same radius as that of the gecko image. This was an important step to ensure that simulated phenotype matches contained, for example, the same number of spots when other statistics such as the fractional area and eccentricity matched. For these disk-shaped regions cut from simulated larger domains, the fractional area and eccentricity of the spots was measured using the same automated scripts as for the live gecko images.

While the LALI-type determines pattern characteristics such as the average fractional area and eccentricity, the domain size of the pattern is relevant because this determines the amount of the pattern that is captured (actual number of spots). A disk-shaped post-orbital head region with domain size that is relatively small compared to the pattern wavelength will have relatively few spots and will be a relatively small sample of the pattern.

### C. Generating the LALI-type to phenotype map by identifying the neutral region of each phenotype and the phenotype cloud of points in LALI-space

To investigate how a produced pattern (the phenotype) depends on the input parameters used for the simulation (the LALI-type), we consider the concept of the LALI-type to phenotype map, or *L*→*P* map (Table 1). More formally, a computational model for a LALI mechanism capable of describing a range of patterns of interest can be defined with a set of model parameters *λ* _1_,*λ*_2_, …, *λ* _*n*_ and a set of rules for evolving the system to generate a pattern. The resulting pattern can be described with a set of morphometric measurements *ρ*_1_, ρ_2_,…,*ρ*_*m*_. We consider that the vector (*λ*_1_, *λ*_2_, …, *λ*_*n*_) is a point in LALI-space (referred as the “LALI-type”) and that the vector (*ρ*_1_, *ρ*_2_, …, *ρ*_*m*_) is a point in phenotype-space (the phenotype) so that the computational model represents a mapping from LALI-space L to phenotype-space P.

Due to the element of random variation in the pattern generation process, a single LALI-type can generate different phenotypes, the ‘phenotype cloud’ of that LALI-type. Likewise, a particular phenotype can be produced by different LALI-types – the region of LALI-space containing this set of LALI-types is the ‘neutral’ region of that phenotype. We describe the LALI-type to phenotype map with the following steps (schematically summarized in Figure 4):

**Figure 4:**
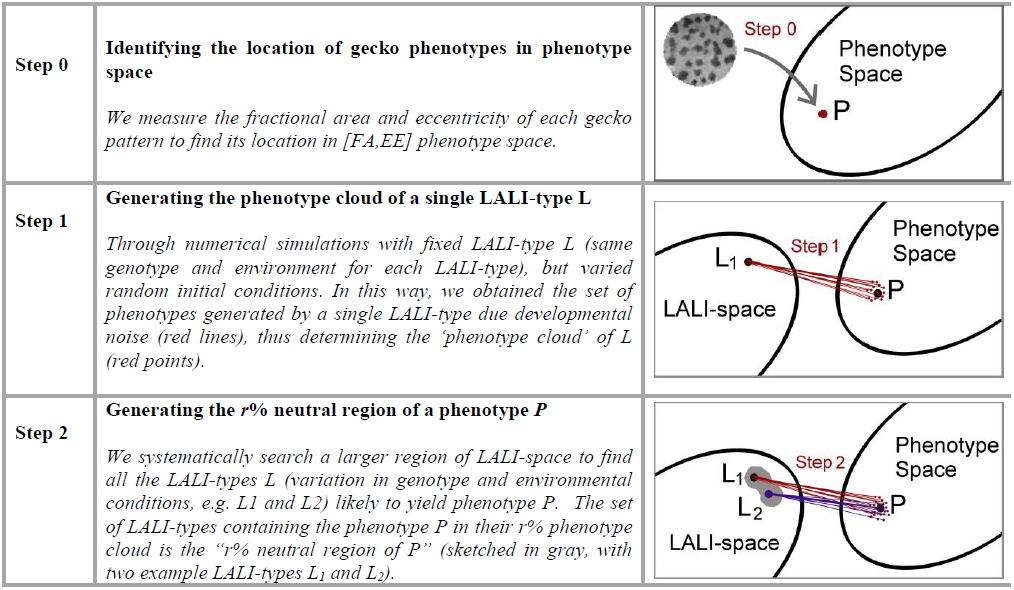
Overview of Methods (Steps 0 – 2) The *L*→*P* map is modeled by one of two computational reaction diffusion models. Due to developmental noise, one LALI-type probabilistically maps to many phenotypes (the ‘phenotype cloud’) and many LALI-types map to one phenotype (the ‘neutral region’ of the phenotype).

#### Step 0

We identify the phenotypes of the eight live gecko patterns, for a small set of selected morphometric measurements, as described above. These phenotypes are points in phenotype space.

#### Step 1: Generating Phenotype Clouds and Defining the Set of Likely Phenotypes

For a given LALI-type, its phenotype cloud is the corresponding distribution of measurements in phenotype space generated from the patterns simulated for that LALI-type. The size of the phenotype cloud describes the role of random variation (developmental noise) for fixed LALI-type. We define the ‘center’ of a phenotype cloud as 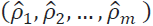 where 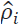 is the average value of the *i^th^* morphometric measure. The center of the cloud and the probabilistic distribution of phenotypes within the cloud are used to define the set of ‘likely’ phenotypes: a phenotype is *likely* if the distance from the phenotype and the center of the phenotype cloud is closer than a specified fraction of the phenotypes in the phenotype cloud. With a pre-specified cut-off, this eliminates phenotype outliers that occur with smaller probability. We define the “ r % phenotype cloud” as the subset of the total phenotype cloud that includes the r% of the phenotypes which are closest to the center. So, points that lie within the 50% phenotype cloud are those phenotypes whose distance from the center is less than the median distance.

#### Step 2: Generating Neutral Regions of a Phenotype

We identify neutral regions of each phenotype in LALI-space. This is describing the way the *L*→*P* map maps *from* regions in LALI-space.

Given a specific pattern phenotype, the *neutral region* of that phenotype is the set of parameters in LALI-space that are “likely” to yield patterns with that given phenotype (using the metric provided from step 1). To systematically find the neutral region of each phenotype, we first identified a region of LALI-space that is capable of producing all eight phenotypes. Conveniently, we achieved this within a relatively low two-dimensional projection for both LALI models. The r % neutral region of a point *P* in phenotype space is then defined as the set of LALI-types *L* for which *P* lies within the r % phenotype cloud of *L*. The larger the parameter *r*, the larger the size of the neutral region.

## 3. Results

### A. Statistical Image Analysis of the Eight Live Gecko Patterns

Figure 5 shows how the phenotype measures varied for each gecko: the fractional pigmented area *FA* of the spots in each pattern (Panel A), the average eccentricity *EE* of the spots in each pattern (Panel B), the average spot size *S* in centimeters for each pattern (Panel C) and the pattern wavelengths (Panel D). The error bars show the dependence of the statistic on the pigment threshold that was chosen to decide if a gray pixel was either white or black. For our analyses (except for the Fourier wavelength), the single threshold of T.= (*μ*.- *σ*) was applied, generating one binary image per gecko photo, so that the value indicated by each filled dot in Figure 5 is the actual value of morphometric parameters in each of the eight images that were used. The spot size was especially sensitive to the threshold that was chosen to binarize the image (relatively large error bars), whereas the measures of the wavelengths of the patterns (average distances between pigmented regions) were robust to this parameter. That is, the size of spots would be larger or smaller depending on whether pixels of intermediate intensity located at the edge of a blotch were classified as pigmented or not. The peak length and Fourier method gave similar measures of the wavelength, however, we speculate that systematic differences might be due to the way the methods average over wavelength variation across the image. For example, based on the way they are calculated, the Fourier wavelength would be biased towards the larger wavelengths whereas the peak length method would be biased towards the shorter wavelengths. The skeletonizations used for calculation of the peak length method are shown in Figure 3, column 3.

**Figure 5:**
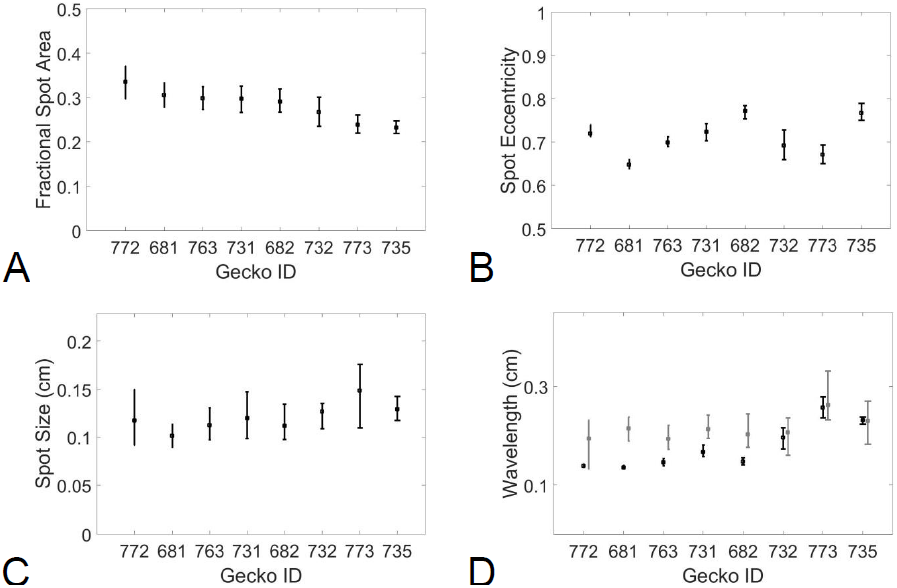
Spot statistics for each Gecko ID. For each binarized image of spot patterning, A) fractional spot area, B) mean spot eccentricity, C) mean spot size and D) wavelength calculated by peak length (black) and Fourier (gray) methods were calculated. Geckos are ordered by decreasing fractional area. Error bars show the minimum and maximum measures of these measures as the threshold for binarization was varied by 0.1σ _i_ where σ _i_ is the standard deviation of the image pixel intensity

#### Step 0: Identifying the Location of the Eight Gecko Patterns in Phenotype Space

For the spot patterns selected from eight live geckos, a set of automated scripts calculated the fractional area (FA) and the eccentricity (EE) of the pigmented spots. The selected *phenotypes* of the eight live geckos patterns, defined as the [fractional area, eccentricity]-pair [FA, EE] located the eight gecko patterns in a two-dimensional [FA, EE]-phenotype-space (‘Step 0’, see Figure 4). The locations of the eight gecko patterns in [FA, EE]-phenotype-space is shown in Figure 6. For the eight live gecko patterns, the fractional area varied from 0.23 to 0.34, while the eccentricity varied from 0.65 to 0.77.

**Figure 6:**
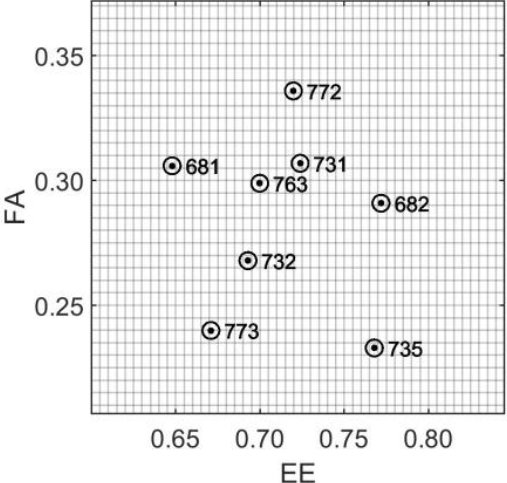
Location of the Eight Gecko Patterns in FA-EE Phenotype Space. The distribution of the eight patterns in [FA, EE] phenotype space where FA is the fractional area of spots and EE is the average eccentricity of the spots.

### B. Describing the LALI-type to Phenotype Map (via the identification of the Neutral Region and Phenotype Clouds of the *L → p* map)

#### Step 1: Generating the Phenotype Cloud of a LALI-type

To illustrate the variation that would occur for a point in LALI-space due to random variation alone, we generate the phenotype cloud of a fixed point in LALI-space (single LALI-type) by simulating 100 patterns at that location in LALI-space. As an example, Figure 7 shows a phenotype cloud of the linear (Turing) LALI map for the LALI-type 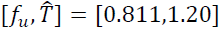 (Panel A) and a phenotype cloud of the non-linear (FitzHugh-Nagumo) LALI map for the LALI-type [*ρ, T*] = [2.63,1.56] of (Panel B). The 100 patterns generated create a cloud in [*FA, EE*]-phenotype space due to stochasticity since the LALI-type, and thus all LALI parameters, were held fixed. These points in LALI-type mapped to a phenotype cloud approximately centered on the phenotype of the gecko pattern #682 (the black point labeled 682 on Panels A and B).

**Figure 7:**
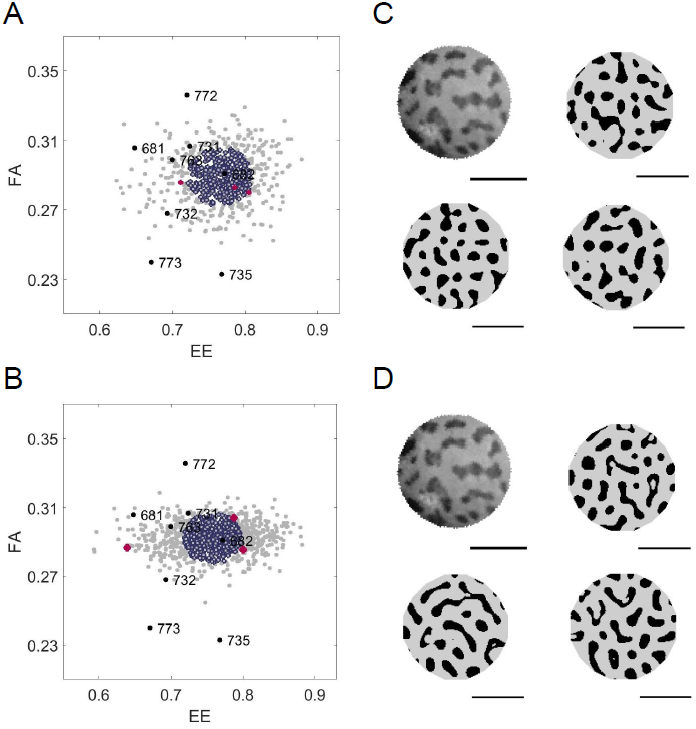
Random Phenotype Variation. The phenotype cloud for 1000 simulations showing the random variation of a single LALI-type, for either the A) linear or B) FitzHugh-Nagumo models. The representative LALI-type was chosen from the 50% neutral region of gecko pattern #682. (The location of this LALI-type is shown as a labeled white dot in Figure 8). The phenotypes of the 1000 simulations are indicated as gray disks in FA-EE phenotype space, while the 500 within the 50% radius of the phenotype cloud are outlined in purple. Three random phenotypes from the cloud are shown in red (see below). C, D) The pattern isolated from the image of Gecko #682 and the patterns of three simulated “clones” (patterns generated with the same LALI-type that is likely to yield pattern #682, but allowing random variation). The result is not necessarily ‘close’ to the pattern #682 (their locations in FA-EE space are indicated in red in the panels ‘A’ and ‘B’.) Horizontal bars indicate 0.5 cm.

In Figure 7, in Panels A and B, the phenotypes of points that are most likely (within the radius of a disk containing 50% of the point distribution) are outlined in purple. The phenotype for the gecko pattern #682 is well within this radius. Thus, the chosen LALI-types are points that would potentially generate gecko pattern #682. In Figure 7, Panels C and D, gecko pattern #682 is shown with three patterns that were randomly selected from within the phenotype clouds of each LALI-type. Within the context of our modeling framework, the variations among these patterns correspond to the random variation of genetic and environmental clones.

#### Step 2: Identification of the Neutral Region for Each Phenotype

For each of the eight phenotypes identified, we recovered a region in LALI-space that was able to produce all eight phenotypes. This corresponds to a range of genetic and environmental variation that can produce all eight phenotypes. For both models, we found that it was sufficient to only vary two LALI parameters. In principle, we would expect that the addition of other morphometrics, measuring more subtle patterns features, could require the variation of a greater number of LALI parameters for fitting more subtle pattern details. For the linear model, we varied the threshold morphogen value T and the activation rate *f*_*u*_. For the FitzHugh-Nagumo model, we varied the threshold morphogen value T and the model parameter ρ. For example, for the linear model, the identified region in LALI-space able to produce all eight phenotypes was a region in which the activation rate varied from 0.80 to 0.84 and the threshold morphogen value relative to the mean morphogen value 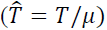 varied from 1.1 to 1.7. The 50% neutral region of each phenotype was identified by searching systematically within this region of LALI-space. Here a LALI-type lies in the 50% neutral region of a phenotype if the phenotype is closer to the center of the phenotype cloud than 50% of the points in this cloud. This means that the phenotype is closer to the ‘typical’ pattern than at least half of the possible patterns. Thus the 50% neutral region of a phenotype can be thought of as those LALI-types that yield a pattern similar to that of the phenotype with high probability. The 50% neutral regions of eight live gecko phenotypes are shown in Figure 8. We found that the 50% neutral regions were all non-overlapping with each other. In principle, the neutral regions might have overlapped if two geckos in the cohort had especially similar patterning. The 50% neutral regions of the phenotypes corresponding to the geckos labeled #763 and #731 nearly overlapped, but shared no grid points in common. The regions of LALI-space shown in Figure 8 (Panel A and B) were extended to show more of the neutral regions of #735 and #682 but these are still cut off in the panels because the neutral regions of #735 and #682 are so elongated. The relative elongation of the neutral regions of #735 whereas in other regions of LALI-space the pattern is more sensitive to small changes in these parameters.

**Figure 8:**
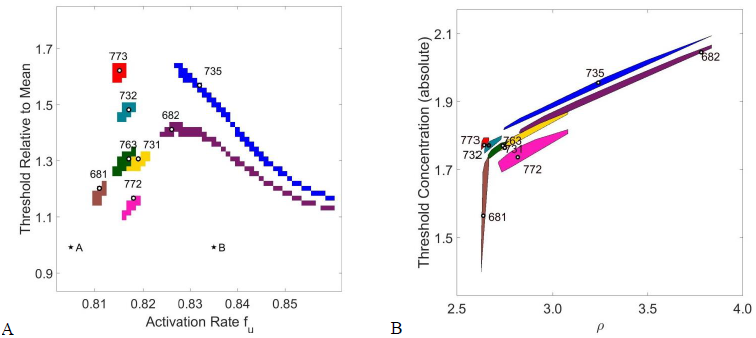
50% Neutral Regions of each of the Eight Gecko Patterns in LALI-space. For each of the eight gecko patterns, we identified the neutral region A) in 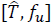-LALI-space for the linear Turing implementation or B) in [*T, ρ*] LALI-space for the non-linear FitzHugh-Nagumo model. The white circle in each neutral region shows the LALI-type chosen to generate representative phenotype clouds in Figures 7 and Figure 8. The labeled stars A and B are the points in LALI-space that are used to generate the “*preternatural*” patterns in Figure 12. (Both models used the *50*% phenotype cloud to generate the 50% neutral region, see the description under “Step 2” in the Methods, section C.)

### C. The LALI-type to Phenotype Map Enhances Understanding of Intra-Group Variation

For both implementations of the LALI-type to phenotype map, the linear Turing model and the FitzHugh-Nagumo model, we show the mapping of a regular grid in LALI-space to the corresponding region in phenotype space. Here a point in LALI-space is mapped to the coordinates of the center of its phenotype cloud in FA-EE-space. The regular grid lines in LALI-space are lines along which one parameter in LALI-space is held constant. For example, the horizontal grid lines in Panel A of Figure 9 (red lines) are lines along which 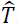 is held constant and these map to non-linear curves in FA-EE-space (red curves in Panel B). Along each of these red curves in Panel B, the value of T is held constant and each curve shows the effect of varying the parameter *f*_*u*_. The extent to which these curves deviate from a pattern of parallel evenly spaced lines shows the extent to which there is 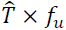 “interaction” (in the sense of statistical interaction, where interaction is the extent to which the effect of the parameters is not additive [46]). Regions of the map where these curves deviate from parallel straight lines are regions where there is more interaction. As an example of one way to interpret these isoclines, if one LALI parameter was genetically controlled and one parameter was environmentally controlled, this is the shape that the G×E interaction would take (see [47]). Both computational models predict that the T-isoclines (red-lines) particularly deviate from straight, parallel lines when the eccentricity is high and the fractional area is low (lower right corner of the grid). Phenotypes in this area of phenotype space (#682 and #735) would be most differentially affected by small changes in the LALI-parameters (corresponding to genotype and environmental variation).

**Figure 9:**
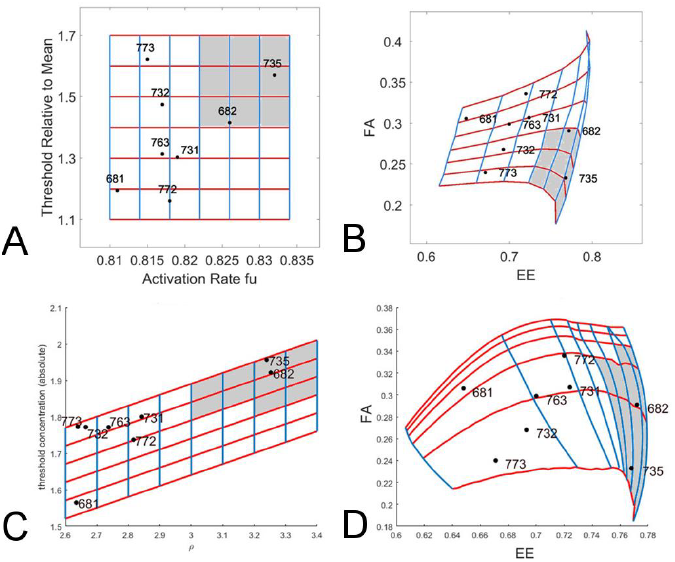
Bias of the LALI-space to phenotype space mapping. A regular grid containing points from the neutral regions of the eight gecko patterns is chosen and the mapping of that grid to phenotype space is shown. A rectangular grid A) in 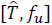-LALI-space for the linear Turing implementation or C) in [*T, ρ*] LALI-space for the non-linear FitzHugh-Nagumo model and the mapping of that grid in FA-EE-space for the B) linear Turing map and D) the non-linear FitzHugh-Nagumo model. The gray region indicates 25% of the area in LALI-space, which maps to a smaller fractional area in phenotype space. This corresponds to a higher likelihood of points (a higher density) in that region of phenotype space.

**Figure 10:**
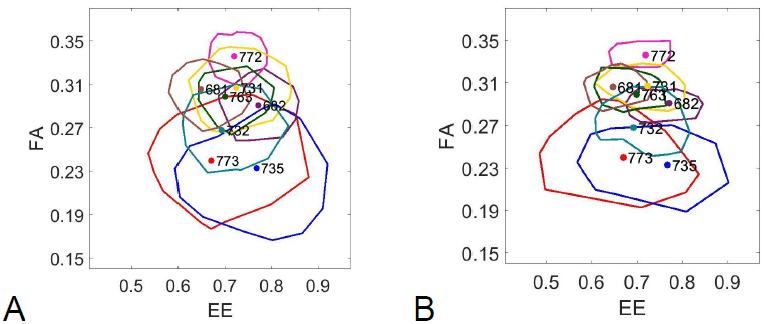
The Intra-Group Variation of the Eight Leopard Gecko Pattern is Larger than Random Variation. Each closed curve shows the outer contour of the 95% phenotype cloud for eight LALI-types that were selected from within the neutral region of each leopard gecko pattern for the A) linear Turing and B) FitzHugh-Nagumo models. These LALI-types are indicated in LALI-space as labeled white dots in Figure 8. Although the phenotype clouds overlap, even the largest phenotype clouds do not contain all of the phenotype variation of the group, indicating that the random variation is not large enough on its own to account for all of the variation.

#### Describing the Bias Introduced by the LALI-type to Phenotype Mapping

There is a mapping bias *for* higher eccentricities and lower fractional areas (Figure 9). To aid in interpreting the bias, we can consider the hypothetical that the parameters randomly vary in the indicated region of LALI-space so that each grid area has an equal probability of being represented by a gecko offspring. However, this does not correspond to equal probabilities of encountering the regions in phenotype space. All the points in the region colored in gray in LALI-space (25% of the total area) map to a relatively small region in phenotype space. This translates to a relatively high probability of points clustering in that region in phenotype space (25% of the points would land in that gray region). In other words, even if the values in LALI-space were randomly varied around a central position, they would map to a phenotype that is skewed towards higher eccentricity and lower fractional areas. Without knowing of this underlying bias introduced by the LALI map, this clustering would be interpreted as a “designed” or purposeful clustering in phenotype space rather than a random one.

Another way of perceiving the bias is that gecko patterns that seem relatively well-separated in phenotype space (e.g., #735 and #682 in Panels C and D) could be found to have a relatively small separation in LALI-space. The relatively small differences in LALI-space result in large differences in phenotype space. Even if one phenotype had a selective advantage over another, it might be difficult to control which phenotype would arise. Also, differences in patterns may be more due to chance than would be expected just by looking at their distances in phenotype space (see also Discussion).

#### Application of the L → p map: Interpreting the Intra-Group Variation in the Context of the Random Variation

Simulations for the same LALI-type yield a range of patterns due to random variation. We investigate and compare the intrinsic variability of each the eight gecko patterns by generating the distribution of patterns for each phenotype that would be expected due to random variation. For each phenotype, we generate a distribution of patterns using a LALI-type selected from their neutral region, and crucially we also restrict the domain size of the generated patterns to match that of the gecko pattern under study (see Methods, Section B). While the LALI-type determines pattern characteristics such as the average fractional area and eccentricity, the domain size of the pattern is relevant because this determines the amount of the pattern that is captured (actual number of spots). A disk-shaped post-orbital head region with domain size that is relatively small compared to the pattern wavelength will have relatively few spots and will be a relatively small sample of the pattern.

Figure 10 shows the outlines of these phenotype clouds for all eight LALI-types, with clouds approximately centered at each phenotype. Here the outline of the 95% phenotype cloud is shown for both the linear and FitzHugh-Nagumo models, see the description under “Step 2” in section 2C above. Due to both random variation and the restricted sample size of the number of spots on the individual gecko heads, there is extensive overlap of phenotype clouds and several LALI-types are capable of producing more than one of the eight gecko phenotypes. The potential of one LALI-type to produce more than one of the observed live gecko phenotype is shown by the inclusion of more than one phenotype in the outline of a phenotype cloud. For example, the LALI-type producing the cloud centered at gecko pattern #735 is also capable of producing a phenotype like that of gecko pattern #773 or even gecko pattern #732 (with a lower probability) (Figure 10). Although there is extensive overlap among the phenotype clouds, no single phenotype cloud includes all of the eight gecko patterns. Thus the variation between the gecko patterns is larger than that of random variation, even when the domain size of the pattern is considered. By defining the radius containing a specified percentage of each phenotype cloud, one can determine the relative likelihood that a phenotype could be generated by a single LALI-type due to random variation. We observe that LALI-types mapping to phenotype clouds in different regions of phenotype space have phenotype clouds with different sizes, corresponding to different amount of random variation intrinsic to that set of LALI parameters.

A domain size that is relatively small compared to the pattern wavelength will have relatively few spots; this is a relatively small sample of the pattern and the intrinsic variability of that phenotype will larger. For example, gecko patterns #735 and #773 are both especially small and have an especially large distance between spots, so that the patterns contain very few spots. Due to the effects of relatively small domain size, these phenotype specimens have the largest intrinsic phenotype variability.

The inclusive relationships of the phenotype clouds in Figure 10 also allow for a description of the estimated likelihood that two gecko phenotypes correspond to the same LALI-type. The corresponding relatedness of pairs of phenotypes is given in Figure 11. Here we determined for each pair (*i,j*) of geckos whether *i* is contained in the 95% phenotype cloud of *j* or vice versa for each of the two models. If this was not the case for any combination, we assigned a score of 0. If this was the case for one model and in one instance, we assigned a score of 1, etc., up to a maximum possible score of 4. In Figure 11, the maximum score is represented by a black field at position (*i,j*), whereas the minimum score 0 corresponds to a white field, with graded gray levels indicating intermediate scores. The two computational models are largely consistent with respect to their predictions regarding the relatedness of the phenotypes (for example, both models indicate it is highly unlikely that a LALI-type producing a phenotype for #772 would also produce a phenotype for #735). The use of graded gray levels in Figure 11 is a way to summarize where the models were not entirely consistent (all pairs that are not white not black) and present the degree of relatedness by weighting the two models equally.

**Figure 11:**
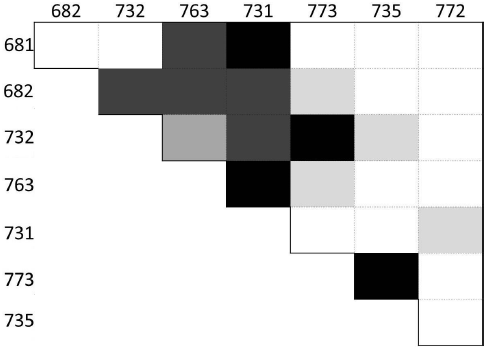
Classification of the relatedness of pairs of phenotypes. Pairs of the geckos IDs 681, 682, 732, 763, 731, 773, 735, 772 are classified according to a measure of relatedness based on the linear and FitzHugh-Nagumo models used in this paper. The main idea of this measure is whether a likely combination of genotype and environmental factors for the head patterning of one of the geckos in a pair can also produce the pattern of the other gecko with developmental noise as the only difference. The darker the color, the closer two patterns are related in this sense, with white color corresponding to the case when neither of the two patterns can be produced by the other’s combination of genetic and environmental factors for any of the models. (See text for the method used to produce the table).

#### Application of the L →P map: generating the likely variation of an observed phenotype

In Figure 7, Panels C and D, gecko pattern #682 is shown with three patterns that were randomly selected from within the phenotype clouds of each LALI-type, showing the typical random variation for a LALI-type within the neutral region of gecko pattern #682. Within the context of our modeling framework, the variations among these patterns correspond to the random variation of genetic and environmental clones. The patterns shown in Figure 3 (columns 4 and 5) represent the closest phenotype matches generated within each phenotype cloud among the 100 simulations for each gecko pattern.

#### Application of the L → p map: generating new patterns outside the likely variation of the group

Just as the *L* → *P* map can be used to generate patterns that are within the expected variation of the group, by choosing points in LALI-space just outside the set of neutral regions generated by the group of leopard gecko patterns, we can simulate patterns beyond the expected variation of the group. Figure 12 shows patterns generated from two LALI-types outside the neutral regions of the other eight leopard geckos (the locations of these LALI-types are shown as A and B starred in Figure 8). This corresponds to patterns generated by environmental and genetic parameters outside the range of variability seen within the gecko cohort.

**Figure 12:**
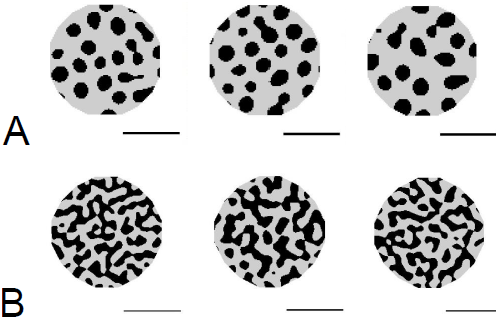
‘Preternatural’ Patterns (Patterns extending beyond the variation observed in the gecko cohort) The LALI framework can be used to generate patterns that are “nearby” in LALI-space, possibly corresponding to the patterns that could be reached by evolutionary change. For each of the ‘starred’ locations in LALI-space indicated in Figure 8, Panel A, we show three random phenotype variations corresponding to that point in LALI-space. These were generated using the linear Turing model. The 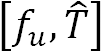 LALI-type for the patterns generated in A and B are [0.805 1.0] and [0.835 1.0], respectively. Horizontal bars indicate 0.5 cm.

## Discussion

Reptile skin patterning has not been as extensively studied as the skin patterning of mammals (felids, giraffes, zebras) and fish (see Introduction for references). However, reptile integuments also frequently display periodic patterning motifs of stripes, spots and mixed ‘labyrinthine’ patterns with extensive individual and species variation, furthering the evidence that LALI patterning mechanisms may be widespread among vertebrates.

### Key result: observed pattern variation of the gecko cohort is more extensive than random variation

In our work, we studied the intra-group variation of patterns selected from a specific region of the head of eight geckos. Considering the variation of the live gecko patterns in [FA, EE]-phenotype space (Figure 6), the phenotypes are distributed roughly uniformly in phenotype space, without showing obvious clusters with a large cluster separation. Clustering of phenotypes would suggest a similar genotype-environment contribution for each cluster. Instead, the observed uniform distribution of phenotypes raises the question whether the observed variation of the eight gecko phenotypes may be due to developmental noise. In other words, the geckos may all have the same genotype and environment background and the observed variation is the random variation resulting from self-organized LALI processes (Figure 2). To address this, we studied the LALI-type to Phenotype (L to P) map for each of the two LALI models we considered. We found that the resulting phenotype clouds for each of the phenotypes did not all overlap (Figure 10) indicating that developmental noise alone cannot explain the variations among the phenotypes. As shown in Figure 11 where each square corresponds to a pairwise comparison of gecko patterns, the lighter the color of a square, the less likely it is that the two phenotypes are generated by developmental noise alone. For example, a LALI-type producing phenotype #732 with an intermediate fractional area is more likely to also result in a phenotype like #731, with a higher fractional area, or #773, with a lower fractional area, but it would be less likely for a LALI-type producing phenotype #773 to also result in a phenotype like #731 as a result of developmental noise alone.

### Matching of observed patterns and selection of morphometric measurements for matching

Using both of the LALI models we considered in this study, we were able to identify LALI-types that generate quantitatively accurate matches to all eight of the observed spot patterns on the heads of eight geckos (Figure 3), where a ‘match’ was defined as producing patterns with the same fractional area and eccentricity. When parameters are varied, periodic LALI patterns vary with respect to the wavelength of the pattern, the size of peaks relative to the wavelength, and a location along a continuum from discrete peaks (spots) to elongated, contiguous peaks (stripes and spirals). For patterns with spots that are well-separated and discrete, relative peak size to peak separation and the elongation of peaks may be quantified with fractional area and eccentricity, respectively. In this work, we were not interested in comparing absolute size differences among the gecko patterns since gecko growth occurs during patterning, resulting in patterns of different size. Since fractional area and eccentricity are dimensionless numbers, and thus scale-independent, they were a natural set of morphometrics for our LALI patterns.

The pattern match observed in our work suggests that a LALI mechanism, in this case, is sufficient to generate the salient aspects of the variation of the color pattern observed on the head of gecko cohort. It is reasonable to consider that a LALI mechanism would work in tandem with other pattern mechanisms, in which case secondary mechanisms could distort the initial LALI pattern and create patterns that are outside the range of LALI parameters space. For example, a LALI mechanism could establish an initial pattern that is then non-randomly stretched by growth along an oriented direction. However, we were able to match the eight observed gecko patterns by varying LALI parameters.

We defined a pattern match as a matching of fractional area and eccentricity that are natural morphometric measures for LALI patterns, as described above. Even though we only used two morphometric measurements in our quantitative analysis to generate our matches, the matches are visually very convincing. However, there were aspects of the patterns that we did not try to match, such as pigment saturation and hue, and the fine-scale texture of the contour of the blotches. These finer details of the pattern would require modeling development at a finer resolution, with more biophysical details such as tissue spatial organization and stages of cell differentiation, which are currently not available to us. On the other hand, there are secondary morphometric measurements, beyond fractional area and eccentricity that could be used to fine tune or further validate pattern matches in the future, such as matching the within-pattern variation of spot sizes, eccentricity etc. While fractional area and eccentricity are aspects of the pattern that we matched in this study, unexpected additional variations among patterns that appear as matches in our results instruct us on salient pattern features and motivate the addition of other variables for fitting. For example, our objective matching algorithm identified patterns as optimized matches even if they included annuli (pigmented regions with a large interior hole), for example, see two annuli in the FitzHugh-Nagumo match for #773 in Figure 3. These annuli occasionally appeared as a best fit match because the algorithm was optimizing only fractional area and eccentricity. This instructs us that if annuli patterns are unwanted, the topology of the pattern should be added as another parameter to match, to exclude regions of parameter space that generate blotches with empty interiors. Rather than apply that topological constraint now, ad hoc, we leave those patterns with annuli as an example of how an objective algorithm for pattern matching avoids subjective bias in pattern selection (if searching by hand, we might have avoided pattern space with annuli) and thus instructs on the most informative features of patterning. It is also worth considering that some variations of leopard gecko patterns may be found with such annuli.

### The L→ P maps provides an important metric for interpreting the distances between patterns

The difference in two phenotypes can be quantitatively described by the difference in the morphometric properties of the phenotypes (head size, tail length, average spot size, etc.). The concept of using a LALI model to index the difference between phenotypes is a compelling idea and was recently applied by Ledesma-Duran *et al.* [48]. Ledesma-Duran and co-workers studied how the range of phenotypes (skin pattern) of *Pseudoplatystoma* fishes could be abstractly quantified (indexed) by the variation of one parameter in their reaction diffusion model. The compelling advantage of using a LALI model for indexing is that seemingly complex phenotypic differences in patterns, such as spots versus stripes versus labyrinthine patterns, can be described by a small number of parameters (for example, as observed in [49]).

Whether the pattern variation is measured by absolute morphometric differences or indexed by a mathematical model, the expected random variation of the pattern provides a necessary context for interpreting the measured difference between two patterns. The geometry of the *L*→ *P* map (that is, how a specific change in LALI-space results in a specific change in phenotype) provides guidance regarding the appropriate way to interpret distance between patterns. For instance, ‘small’ changes in LALI space (due to a combination of genotype and environmental changes) may correspond to large changes in pattern appearances (due to developmental noise). In these cases, the two phenotypes may appear to be very different, when in actuality, they are closely related. The opposite situation may also occur, where two phenotypes may seem similar but they would originate from different regions in LALI-space. Our methods help to look beyond morphometric measurements and uncover such ‘hidden’ relatedness.

Further, details of the geometry of the *L* to *P* mapping may provide clues into the ways that the freedoms and constraints of the developmental mechanism shape the effect of selection pressures on color patterning. The large neutral regions of #735 and #682 indicate that LALI parameters could vary within these regions, with little change in the resulting phenotype, so that parameters could drift in a neutral manner to new regions of LALI-space. Varying sizes of the phenotype clouds indicate that different regions of the *L*→*P* map differ in their intrinsic variability. If there is selection for more or less variation, then this may shift the selection pressure from particular phenotypes to particular regions of LALI-space. A rich literature of theory can be applied to such phenotypic landscapes, such as the neutral theory of Kimura [50, 51], survival of the flattest [52] and arrival of the frequent [53].

### Generalizability of this Framework for Other Models of Developmental Noise

The approach we describe here for generating a metric for the variation of patterning that involves separating the fixed (genotype and environmental) versus stochastic sources of phenotype variation can be applied to any developmental mechanism that permits modeling of pattern generation from a set of fixed initial conditions with stochastic effects.

A local activation long range inhibition (LALI) mechanism is a likely candidate for a patterning mechanism for spot patterning on the heads of a cohort of geckos, especially due to the familiar combination of periodic spots and stripes throughout their body color plan and development, but the molecular details of the mechanism are not known. Rather than commit to a LALI mechanism and yield results that are potentially narrower, we compare two implementations of a LALI core logic: one linear and one non-linear, both reaction diffusion mechanisms. The results of the two LALI implementations are similar overall, especially in the geometry of the maps. This overall similarity can be seen by comparing the relative position and sizes of the neutral regions for each gecko pattern phenotype (Figure 8), the similar direction of the bias of the two maps (Figure 9), and the comparable relative sizes and overlaps of the phenotype clouds (Figures 7, 10). The commonalities of the *L*→ *P* maps for these two LALI models will include common properties of LALI maps in general while the more subtle differences between the models (relative size and detailed shape of the phenotype clouds, and the extent of overlap) are differences that can be expected from one LALI mechanism to another. In future work, a broader range of LALI mechanisms might be compared, or it would be interesting to compare the topology of maps generated by more diverse non-LALI mechanisms. This would be especially interesting if differences in the geometry of variation make testable predictions to distinguish competing mechanisms, as suggested by [39] in the context of assessing the effect of parameter perturbations.

### Further Applications

Here we describe an approach for taking a very small set of patterns (only eight individual gecko patterns were compared in our study) and describing their relative relationship (separation, closeness, potential for overlap) within the context of the LALI framework. Without the underlying LALI framework, a much larger number of pattern specimens would be needed to describe a metric for their variation. That is, when two patterns appear to be similar, measuring the differences between very many patterns would be needed to quantitatively determine whether the difference between these two patterns is relatively small or large, compared to the average variation observed among the pattern specimens. Within the context of the LALI framework, even a single pattern specimen can be identified as a region in LALI space (i.e., the set of LALI parameters specific to a given LALI model that is likely to generate this pattern). Once a location has been identified, the LALI model can be interrogated to determine the amount of stochastic variation that can be expected for that location in LALI space. Within the LALI framework, there is the potential to fruitfully compare the patterns of even a small number of pattern specimens by determining whether their expected ranges of random variation would include one another. There is also the potential for comparisons across species, regardless of differences in the specific underlying LALI mechanism, since patterns are mapped by their morphometric characteristics, not their underlying biophysical parameters. Thus, by mapping species pattern movements in a common LALI space, future investigations may find patterns of variation (for example, long term speciation trends) that are common across species, clades, etc. As we have done here, the LALI framework can be used to determine the fraction of observed diversity that is due to genetic and environment factors, which determines the extent to which specific patterns are inheritable or reproducible. The contribution of developmental noise to pattern diversity can be significant (for example, in [1]) and contribute a selective advantage (‘bet-hedging’ [54]).

Finally, the LALI framework can be used to explore patterns that are just beyond the space apparently explored by the individual variability, to yield ‘supra-natural patterns’ that may or may not be represented in the wild. From an evolutionary and ecological point of view, identifying potential color patterns that do not occur in wild animal populations, but that can instead be generated either mathematically or sometimes by targeted captive-breeding efforts, provide a ground to investigate the evolutionary constraints (e.g., selective, genetic or developmental) that impede the occurrence of these phenotypes in nature.

## Conclusion

With image analysis and the selection of several morphometric features for analysis, a difference between two phenotypes can be measured. Without a model of stochastic variation like that of the *L*→*P* map, however, it is difficult to interpret the significance of a small difference in phenotypes. The model of variation presented here provides a framework for contextualizing the noisy difference between phenotypes within the context of random variation due to developmental noise. Further, the set of parameters in LALI-space that yields a matching phenotype is an efficient summary of the set of real-world parameters that would be needed to specify that phenotype, for any biological mechanism described with a core LALI logic. This points to a method for classification and comparison of patterns across a broad range of contexts. This work underscores the need for significant interdisciplinary effort [8] to advance biometric approaches for generating and analyzing phenotype data.

## Acknowledgments

We are thankful to Julien Claude and Scott Glaberman for comments on an early version of this manuscript. J. Anderson, E. Brumeier, C. Cummings, M. Detjen, I. Eid, and D. Zarkower for help with gecko care and maintenance and I. Eid for photographing geckos.

